# Acoustic indices predict recovery of tropical bird communities for taxonomic and functional composition

**DOI:** 10.1101/2025.03.25.645239

**Authors:** Sonja Kümmet, Jörg Müller, Zuzana Burivalova, Martin H. Schaefer, Rudy Gelis, Juan Freile, Annika Busse, Marcel Püls, Peter Kriegel, Sebastian Seibold, Nico Blüthgen, Maria de la Hoz, Matthias Schleuning, Eike Lena Neuschulz, Oliver Mitesser, Mareike Kortmann

## Abstract

Quantifying the success of biodiversity restoration is a major challenge in the UN Decade on Ecosystem Restoration. We evaluated the potential of acoustic indices to predict the recovery success of bird communities within abandoned agricultural areas. Using audio recordings from a lowland tropical forest region, we identified 334 bird species and calculated established acoustic indices. Community composition was analyzed using Hill numbers, accounting for incomplete sampling. Acoustic indices effectively predicted independent species data (R^2^ = 0.59–0.76), capturing not only taxonomic but also functional and phylogenetic composition. Taxonomic composition was best predicted for common and dominant species, while functional and phylogenetic compositions were more accurately predicted for rare and common species. In addition, community composition was strongly influenced by the surrounding habitat. Our findings demonstrate that a small set of acoustic indices, once validated by stratified ground truth data, provides a powerful tool for assessing restoration success over large tropical areas, even for functional composition of rare tropical birds.

## Introduction

Biodiversity loss ranks among the most pressing global challenges, with profound consequences for ecosystems and human well-being (Díaz et al. 2019; Isbell et al. 2017). The restoration of tropical forests is critical in addressing this crisis, given the unparalleled biodiversity and ecosystem services they provide (Pillay et al. 2022; Hubau et al. 2020). While efforts to restore tree cover have made notable progress, questions remain about the recovery of the diverse flora and fauna that underpin ecosystem functioning (Chazdon et al. 2009). Monitoring this diversity is essential for a realistic and comprehensive assessment of restoration success. This need becomes even more pressing as biodiversity credits come into play as a tool for financing conservation initiatives (Antonelli et al. 2024). To ensure their meaningful application and to prevent misuse, robust methods for assessing conservation success are crucial (Vardon und Lindenmayer 2023). However, traditional biodiversity monitoring methods, based on expert observation and identification of species, are expensive, time-intensive, and poorly scalable (Ford et al. 2024). To overcome these limitations, innovative approaches are needed. Acoustic monitoring has emerged as a promising tool, enabling the survey of a wide range of sound-producing species in a standardized, cost- and time-effective manner (Teixeira et al. 2019; Vega-Hidalgo et al. 2021).

We tested the potential of acoustic indices to predict the recovery success of bird communities in abandoned agricultural areas in the Ecuadorian Chocó Forest. This region is characterized by small-scale agriculture, surrounded by forest or agriculture (Escobar et al. 2025; Gindhart et al. 2024). Within this landscape, we examined a recovery gradient ranging from active pastures and cacao plantations, through regenerating forests to old-growth forests. Based on 2380 audio files, we identified 334 bird species and calculated well-established acoustic indices. These indices were selected to capture key aspects of acoustic environments: sound coverage across the frequency range (Soundscape Saturation), sound diversity over time (Acoustic Complexity Index, Temporal Entropy, Entropy of Frequency) and acoustic activity (Events per Second), all described in (Müller et al. 2023).

Acoustic indices are known to reflect patterns of land use (Burivalova et al. 2018) and forest recovery (Müller et al. 2023). However, previous studies have not accounted for critical aspects of biodiversity quantification, such as sample coverage (Chao und Jost 2012), differences in species frequency distributions by Hill numbers (Hill 1973), and the functional characteristics of bird communities (Chao et al. 2014). While taxonomic diversity typically recovers relatively quickly or remains consistent along restoration gradients (Kortmann et al. 2025), species composition tends to recover at a much slower pace (Dunn 2004; Rozendaal et al. 2019). Consequently, relying solely on taxonomic diversity risks to overestimate the restoration success.

To get a more precise estimate, we calculated community composition using incidence-based frequencies (most suitable for acoustic data) along the Hill numbers, explicitly accounting for incomplete sampling (Kortmann et al. 2025). To evaluate functionality and resilience across different habitat types, we assessed taxonomic, functional and phylogenetic diversity. Given that many traits are phylogenetically conserved, phylogenetic diversity serves as a reliable proxy for ecological differences among species in many taxa (Cadotte et al. 2012; Bregman et al. 2016; Thorn et al. 2018).

This study advances the field by demonstrating the effectiveness of acoustic indices derived from autonomous audio recordings in assessing restoration success. Importantly, it expands on prior research by exploring the potential of acoustic indices to predict communities from independent data and capture three dimensions of diversity (Burivalova et al. 2018; Müller et al. 2023). By including species spanning a gradient of rarity – from infrequent to highly frequent – and examining sites across various stages of regrowth within forested and agricultural landscapes, our research provides critical insights into ecosystem recovery. Together, these findings address a key challenge in conservation: developing consistent, reliable, and credible measures of biodiversity recovery (Antonelli et al. 2024).

## Methods

### Study site

The raw data for our study were collected across 85 plots in Canandé, a biodiversity hotspot located in northwestern Ecuador dominated by lowland tropical rainforest. 65 plots were established by the REASSEMBLY research unit (http://www.reassembly.de) (Map 1).

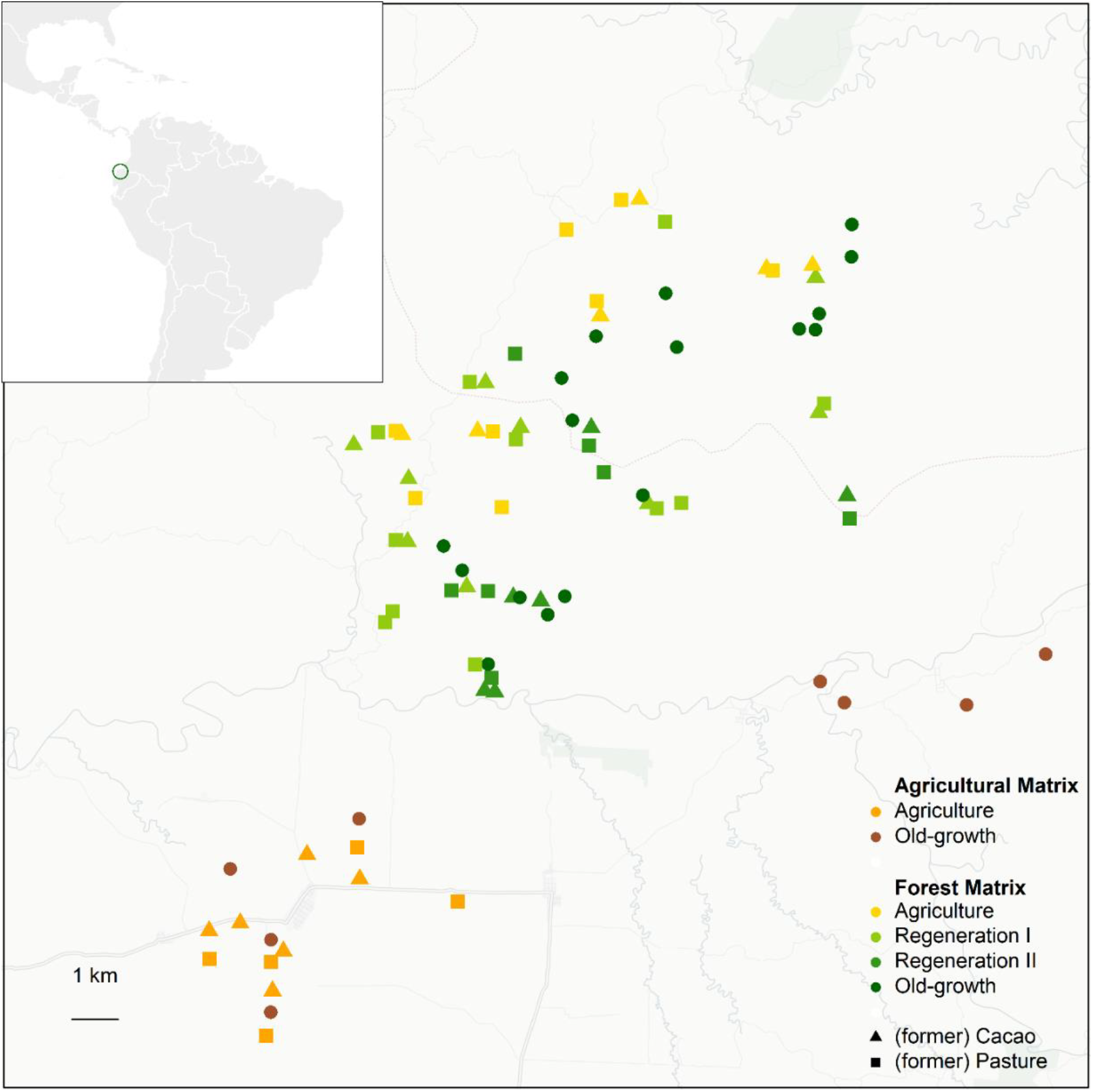

**Map 1** shows the location of the 85 study plots in Canandé, northwestern Ecuador. 65 of the 85 plots are surrounded by forest (“Forest Matrix”), the other 20 by agriculture (“Agricultural Matrix”). The plots span a restoration gradient ranging from active agriculture to secondary forests of regeneration stage I (1–20 years after abandonment), secondary forests of regeneration stage II (21–38 years after abandonment) and old-growth forests.

As they are surrounded by forest, we refer to them as the ‘forest matrix’. The remaining 20 plots, situated in a landscape dominated by agricultural land use, form the ‘agricultural matrix’. On average, forest matrix plots have 78% forest cover within a 1 km radius, compared to 6% for agricultural matrix plots. All 85 plots represent a chronosequence of forest recovery, ranging from actively managed pastures and cacao plantations through restoration areas – early regeneration stages (1–20 years after abandonment) and later regeneration stages (21–38 years after abandonment) – to old-growth forests (Fig. 1). Detailed information about the study area and design can be found in (Müller et al. 2023; Escobar et al. 2025).

**Figure 1.**
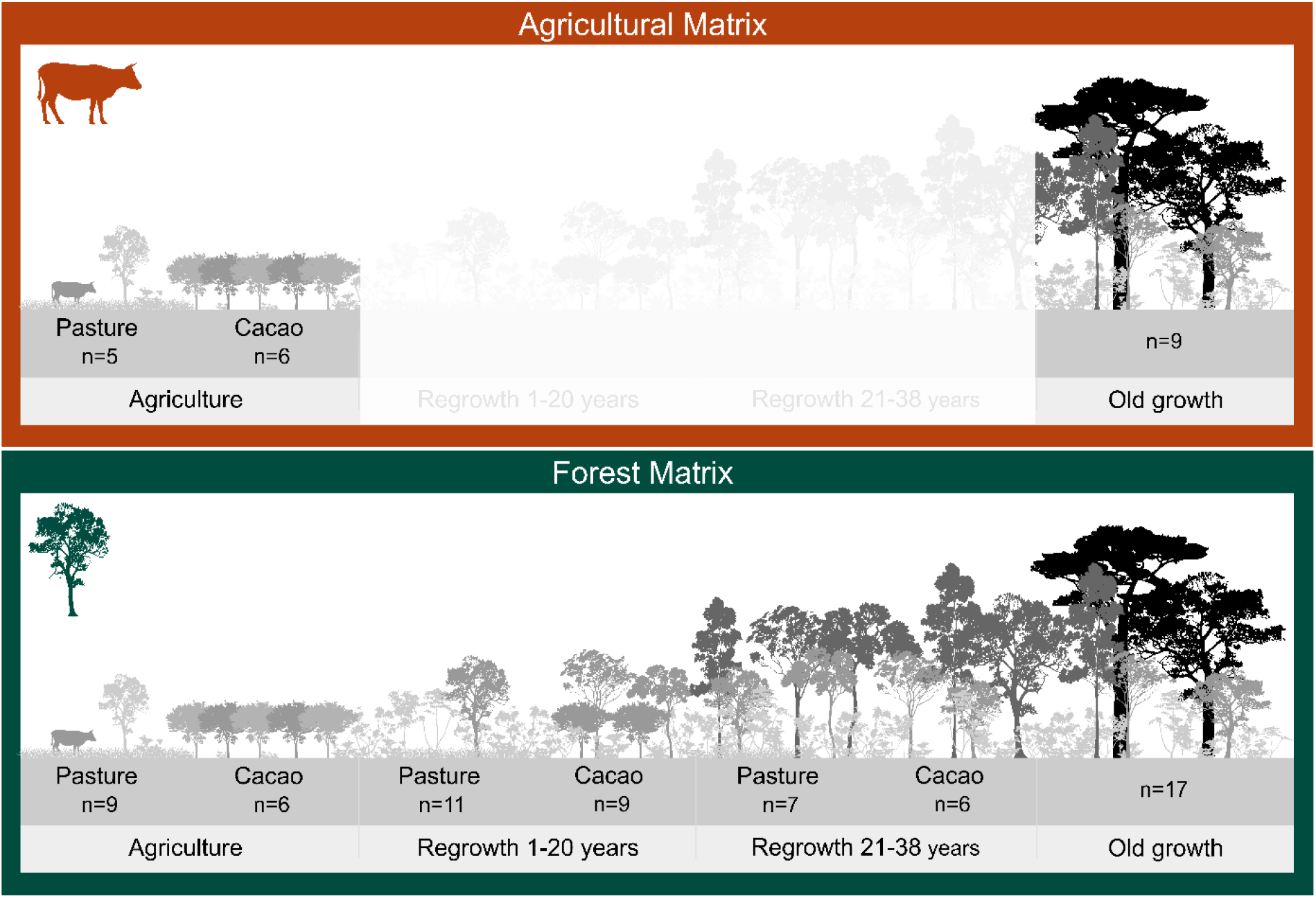
shows the division of the 85 Ecuadorian study plots into the “agricultural matrix” (20 plots surrounded by agriculture) and the “forest matrix” (65 plots surrounded by forest). In the “agricultural matrix,” only the two extremes are present: plots with active land use (pasture and cacao) on one end and old-growth forests on the other, with no recovery stages in between.

## Data Collection

At the center of each plot, approximately 1.70 m above the ground, we placed a bioacoustic recorder (BAR-LT, Frontier Labs, Meanjin, Australia) equipped with a downward-facing omnidirectional microphone. These recorders captured 2-minute .wav files at a sampling rate of 44.1 kHz every 15 minutes throughout the day during two periods: October 12 to November 28, 2021, and October 21 to December 19, 2022.

A subset of the recordings – 28 files per plot – was analyzed by two local expert ornithologists: 14 × 85 files by Juan Freile and 14 × 85 files by Rudy Gelis. The criterion for selecting the files was to capture high bird activity phases around dusk and dawn, as well as flock activity during the day. To achieve this, recordings from 06:00, 06:15, 06:30, 06:45, 07:00, 07:15, 12:00, 12:15,

16:00, 16:15, 17:00, 17:15, 18:00 and 18:15, across two days, were chosen. For each file, the ornithologists determined the presence (1) or absence (0) of 334 bird species. The resulting community composition dataset comprises 2.380 rows (one per file) and includes filename, plot and detection/non-detection for the 334 bird species.

In addition to the expert-based identification of bird species, we used the toolbox *AnalysisProgram.exe* to calculate acoustic indices for another subset of files. For each plot and year, we randomly selected 14 days with at least 96 files per day, except for four plots that had only 13 days meeting this criterion. Acoustic indices were calculated for 60 s segments without overlap based on FFT windows of 512 data points for spectral decomposition. The results were aggregated by calculating the mean value for each plot.

The morphological and ecological bird traits used in this study were obtained from the AVONET database (Tobias et al. 2022). The phylogenetic tree was built with the U.PhyloMaker package in R (Jin und Qian 2023), based on the bird-megatree by (Jetz et al. 2012).

## Analysis

All statistical analyses were conducted with R version 4.4.0 (2024-04-24 ucrt). We used an adapted version of the iNEXT.beta3D function from the *iNEXT.beta3D* package – iNEXTbeta3D_pair3D – to compute pairwise *β*-diversity indices (distance matrices) for all 85 study plots, taking into account three dimensions of biodiversity – “taxonomic diversity” (TD), “phylogenetic diversity” (PD) and “functional diversity” (FD) – and focusing on infrequent (q = 0), frequent (q = 1), and highly frequent (q = 2) species (based on Jaccard, Horn, and Morisita-Horn index). Within the Hill numbers framework, the exponent q determines the weighting of species relative abundances.

Our input data included the community composition dataset (datatype = “incidence_raw”), a phylogenetic tree for PD (in Newick format) and a species pairwise distance matrix (Gower distance, computed from the traits) for functional diversity (FD).

Kortmann et al (2025) shows that sample coverage significantly declines along the recovery gradient in our study area, even with consistent and highly standardized sampling effort. To address this bias, we standardized sampling coverage by using the mean of the 85 sample coverages estimates (SC(2n)), as provided by the DataInfobeta3D function (Supplementary Figure S1).

To reduce the multidimensional complexity of the distance matrices to a two-dimensional representation, we performed an ordination with each distance matrix (TD, PD and FD) and for all orders of q (q = 0, q = 1 and q = 2) using the metaMDS function from the vegan package (Dixon 2003). For two cases (TD q = 0 and PD q = 0), the stress levels exceeded 0.2, indicating that the community data were too complex to be adequately represented in two dimensions. However, for consistency and comparability, we retained the two-dimensional representation.

The resulting nmds axis 1 values were then taken as response variables in linear models, with five acoustic indices as predictor variables. We selected the same set of acoustic indices – Soundscape Saturation, Entropy Of Variance Spectrum, Acoustic Complexity, Temporal Entropy and Events per Second – as in (Müller 2023), due to their demonstrated effectiveness. To move beyond correlation and assess predictive power, we trained the models on two-thirds of the data and reserved one-third for prediction. The split into training and testing data was stratified by NMDS axis 1 – by first sorting the dataframe by NMDS axis 1 and then selecting every third row for the prediction set. This approach maintained a balanced representation of community composition across both subsets.

## Results

The first axis of the ordination of the bird communities consistently reflects the recovery gradient, regardless of the diversity type (TD, PD, FD) or Hill number (q = 0, q = 1, q = 2) (Figure 2 and Supplementary Figure S2). From left to right, agricultural sites (cacao and pasture) are followed sequentially by early regeneration stages (1–20 years after abandonment), later regeneration stages (21–38 years after abandonment), and old-growth forests. All plots within the ‘agricultural matrix’ are shifted along the first axis towards active agricultural patches: The cacao and pasture plots of the ‘agricultural matrix’ are on the far left, while the old growth-forest sites are similar to the early regeneration plots within the “forest matrix” – demonstrating a double effect of agriculture at the plot and landscape scale. Beyond these general trends, the extent and overlap of the community polygons, which capture both the distribution and similarity of communities, vary across diversity types and Hill numbers (Supplementary Figure S2).

**Figure 2.**
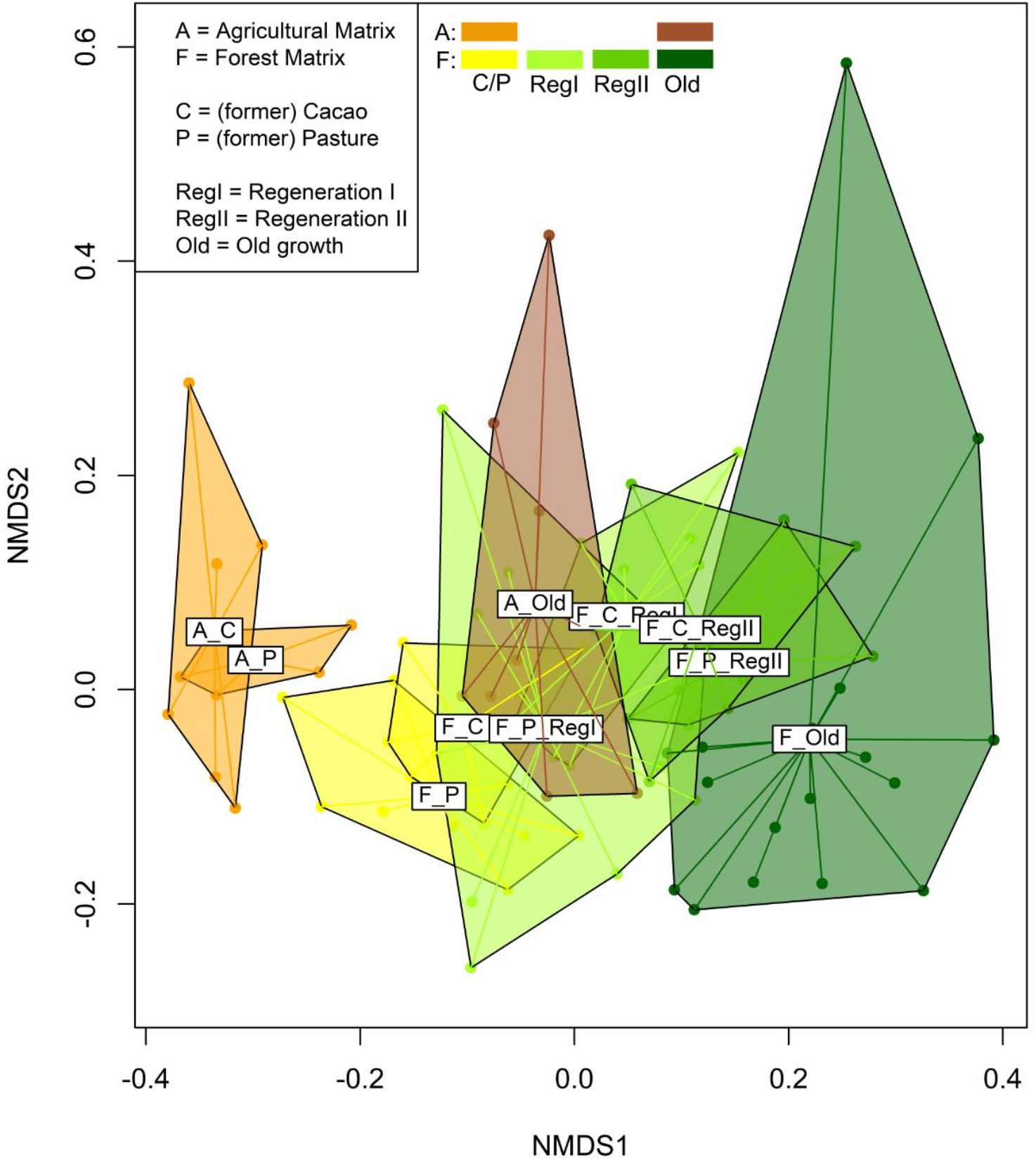
plots the first two nmds axes (NMDS1, NMDS2) of Ecuadorian bird communities, exemplary for Phylogenetic Diversity, q = 0. The category labels consist of up to three components: first, the matrix type (“A” for Agricultural matrix, “F” for Forest matrix), second, the type of (previous) land use (“C” for cacao plantation, “P” for pasture) and third, the regeneration level: “RegI” for 1-20 years since active use, and “RegII” for 21-38 years of regeneration. “Old” stands for old-growths forests. The color palette of the “forest matrix” plots ranges from yellow to dark green, the “agricultural matrix” plots are colored in shades of orange and brown.

There were notable differences among the three diversity types (taxonomic, phylogenetic and functional) and orders of q (infrequent, frequent and highly frequent species): For taxonomic diversity, the predictive power, indicated by the R^2^ value, is relatively high (Figure 3). The strongest predictive relationships are observed for frequent and highly frequent species, with a slightly lower value for infrequent species. For functional diversity, which captures the range of functional traits within a community, R^2^ is higher for infrequent and frequent species and lower for highly frequent species. A similar pattern emerges for phylogenetic diversity, which reflects the evolutionary relationships among species. The R^2^ values are higher for infrequent and frequent species and lower for highly frequent species, mirroring the trend seen in functional diversity.

**Figure 3.**
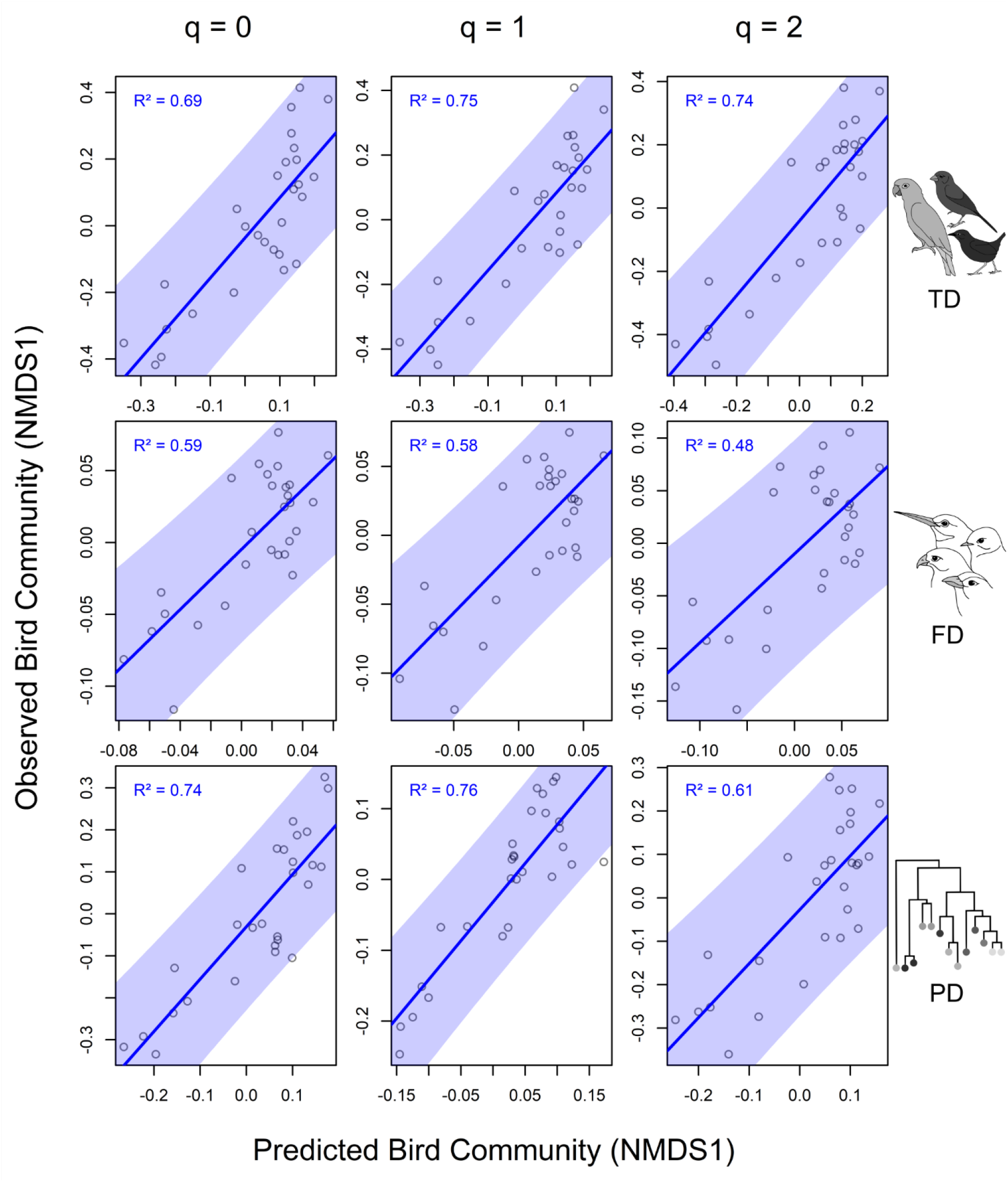
displays scatterplots of the observed NMDS axis 1 values, as proxy for the sampled Ecuadorian bird community, on the y-axis, against the predicted NMDS axis 1 values on the x-axis. The plots are arranged by diversity type (rows: Taxonomic, Functional, Phylogenetic) and orders of q (columns: q = 0 (infrequent), q = 1 (frequent), q = 2 (highly frequent). The predictions are based on a combination of five acoustic indices: Soundscape Saturation, Entropy of Variance Spectrum, Acoustic Complexity, Temporal Entropy, and Events per Second. The blue-shaded 95% prediction interval indicates the range in which future observations are likely to fall.

The impacts of the five acoustic indices on the prediction of NMDS axis 1 vary across the three diversity types and orders of q. Corresponding to the R^2^ values, the effects of the indices are more pronounced for q = 1 and q = 2 in taxonomic diversity, and for q = 0 and q = 1 in functional and phylogenetic diversity. High t-values are particularly evident in frequent and highly frequent species for taxonomic diversity, and in infrequent and frequent species for phylogenetic diversity (Table 1). Despite these variations, the direction of the effects remains largely consistent across all diversity types and orders of q: Soundscape Saturation and Entropy of Variance Spectrum generally show positive effects, whereas Acoustic Complexity and Events per Second show strong negative effects. Temporal Entropy predominantly has positive effects, with two notable outliers for functional diversity at q = 1 and q = 2.

**Table 1.** summarizes the t-values of the five acoustic indices (Soundscape Saturation, Entropy of Variance Spectrum, Acoustic Complexity, Temporal Entropy, and Events per Second) for predicting NMDS axis 1 across the three diversity types (Taxonomic, Functional and Phylogenetic) and orders of q (q = 0, q = 1, q = 2).

| Acoustic Indices | t-values |  |  |  |  |  |  |  |  |
| --- | --- | --- | --- | --- | --- | --- | --- | --- | --- |
|  | Taxonomic Diversity |  |  | Functional Diversity |  |  | Phylogenetic Diversity |  |  |
|  | q = 0 | q = 1 | q = 2 | q = 0 | q = 1 | q = 2 | q = 0 | q = 1 | q = 2 |
| Soundscape Saturation | 1.39 | 1.81 | 2.20 | 0.78 | 1.05 | 0.89 | 1.62 | 1.93 | 1.75 |
| Entropy Of Variance Spectrum | 1.74 | 2.00 | 2.11 | 2.03 | 1.48 | 1.04 | 1.78 | 2.93 | 1.46 |
| Acoustic Complexity | -3.88 | -4.13 | -4.18 | -3.58 | -3.67 | -3.43 | -3.72 | -3.40 | -2.74 |
| Temporal Entropy | 1.15 | 1.23 | 1.11 | 0.50 | -0.18 | -0.70 | 1.77 | 2.61 | 0.55 |
| Events Per Second | -3.15 | -3.54 | -3.72 | -3.68 | -2.86 | -2.28 | -3.53 | -2.86 | -3.30 |

## Discussion

### Three-dimensional Evidence of Restoration Success

Our analysis of bird community composition, based on sound data collected in northern Ecuador, revealed that recovery success is evident not only in taxonomic diversity but also in functional and phylogenetic diversity (Figure 2) – dimensions that are typically overlooked in restoration assessments (but see (Edwards et al. 2017; Hughes et al. 2020)). Additionally, the community composition in old-growth and agricultural patches within the agricultural matrix diverged further from their equivalents within the forest matrix (Figure 1). This suggests that protecting old-growth forests is more effective within a forest matrix compared to an agricultural matrix, underscoring the profound impact of the surrounding landscape on species diversity and community composition (Prevedello und Vieira 2010). Examining the three measures of biodiversity (taxonomic, functional, phylogenetic) together provides deeper insights into ecological functionality and evolutionary relationships, offering a more comprehensive and realistic measure of restoration progress (Kortmann et al. 2025; Rozendaal et al. 2019). This holistic perspective is especially critical for assessing complex and dynamic ecosystems, such as regenerating tropical forests.

### Effectiveness of Acoustic Indices

Expert-based sound analysis is costly, time-consuming, and limited in scope. To overcome these limitations, we tested the predictive power of acoustic indices as an alternative monitoring approach.

Our results (Figure 3) are promising: a combination of five acoustic indices—Soundscape Saturation, Entropy of Variance Spectrum, Acoustic Complexity, Temporal Entropy, and Events per Second—achieved relatively high R^2^ values, ranging from 0.59 to 0.76, for all three biodiversity dimensions: taxonomic, functional, and phylogenetic diversity. Notably, our findings reveal an exciting pattern: while taxonomic diversity showed the strongest predictive power for frequent and highly frequent species, functional and phylogenetic diversity were best predicted for frequent and infrequent species. These results highlight the potential of acoustic indices to predict not only more vocal species but also less vocal or cryptic and rare ones. In our study, infrequent species included those with low vocal activity and those with true rarity. The latter are often key indicators of ecosystem functioning and stability due to their specific habitat requirements (Mouillot et al. 2013).

Previous studies have highlighted the potential of acoustic indices in restoration contexts (Müller et al. 2023; Borker et al. 2020; Lamont et al. 2022). Our findings reinforce this promise by demonstrating that acoustic indices possess strong predictive power. They effectively model the mixed effects of landscape and local habitat on local bird communities and capture complex aspects of species communities, including taxonomic, functional, and phylogenetic diversity, for both dominant and rare species.

Examination of the individual acoustic indices showed that their performance varied across the different biodiversity dimensions, though no consistent patterns emerged. For instance, Temporal Entropy generally performed worse than the other indices but showed a strong correlation with the phylogenetic diversity of frequent species (Supplementary Figure S3).

Temporal Entropy mainly reflects temporal variability in sounds (Flowers et al. 2021), meaning low Temporal Entropy values can be correlated with constant noise, like in urban landscapes. Hence between natural landscapes, the differences should be rather low. In contrast, Acoustic Complexity exhibited the strongest overall correlations with all diversity measures. Acoustic Complexity is a very popular index, used in many ecological studies (Farina 2025). It also showed clear differences between land-use types in a study in Madagascar (Dröge et al. 2021). The complementary nature of the indices highlights the suitability of this selection for capturing variation across different diversity dimensions. This aligns with previous research demonstrating the effectiveness of these indices (Alcocer et al. 2022; Müller et al. 2023).

### Applicability for Monitoring Restoration Success

As acoustic indices remain somewhat cryptic, evaluating their predictive power is essential in biodiversity monitoring, particularly for conservation managers. However, due to the relative consistency of species communities in areas with similar ecological conditions — such as the lowland Chocó forests, spanning from Ecuador to Panama — only a small, stratified subset of data needs to be identified to species level to train predictive acoustic indices models. Once such a training dataset is established, large-scale acoustic recordings can be used to monitor conservation success more efficiently and cost-effectively (Teixeira et al. 2019).

It is crucial, however, to acknowledge the limited transferability of acoustic indices across regions and taxa. Different taxa respond variably to environmental changes (e.g. amphibians and reptiles (Thompson und Donnelly 2018)), meaning the applicability of specific indices cannot be universally assumed. New scenarios require new training datasets and validations to ensure robust and accurate predictions.

### Acoustic Indices versus AI

Our finding that bird communities provide an excellent reflection of local recovery gradients, and even account for the surrounding landscape, naturally raises the question: Why use acoustic indices as a proxy instead of directly identifying species communities with AI? Given the rapid advancements in AI technologies, including CNN (Convolutional Neural Network) models like Merlin and birdNET (Xie et al. 2023), this is a valid consideration.

However, the primary limitation lies in the availability of labeled training data, particularly for rare species. Current AI models often capture only a fraction of the community due to the lack of comprehensive datasets. Extensive datasets are typically available for well-studied areas such as the US and Europe (Gibb et al. 2019). In contrast, libraries for hyperdiverse regions, where biodiversity is both exceptionally high and critically endangered, remain incomplete (Zamora-Gutierrez et al. 2016). This gap is particularly pronounced in tropical rainforests, as highlighted in previous studies (Müller et al. 2023; Sun et al. 2022).

### Acoustic indices for Biodiversity Credits

The search for efficient biodiversity monitoring tools is closely tied to the global debate about establishing a compensation system for biodiversity, akin to carbon credits (Antonelli et al. 2024). While tree biomass recovery can be efficiently tracked through satellite data (Le Toan et al. 2011), monitoring the recovery of tree species requires much more detailed and ground-based information (Asner et al. 2014). Assessing animal biodiversity recovery is even more complex, especially for bird communities that span a vast range of diversity: from tiny hummingbirds to the majestic Harpy Eagle, from silent species like the Banded Ground Cuckoo to loud ones like the Great Tinamou, and from conspicuous species like Toucans to cryptic nocturnal birds like Nightjars.

Our results demonstrate that a small set of acoustic indices can effectively describe this immense diversity in terms of species, evolutionary lineages, and traits. According to the SAGED criteria outlined by Ford (Ford et al. 2024), which define essential requirements for biodiversity monitoring tools in the context of biodiversity credits, acoustic indices meet the standards for being *Scalable, Accessible* (in terms of cost and expertise), and *Evidenceable*. Only in terms of *Direct measurability* and *Granularity* are acoustic indices outperformed by other methods. However, our findings suggest that integrating acoustic indices with expert-based community data can merge the strengths of both approaches, yielding a monitoring method that satisfies all SAGED requirements.

By providing a robust, scalable framework for assessing multiple dimensions of animal diversity, we establish a critical baseline for biodiversity credits. This approach is particularly valuable in hyperdiverse tropical forest regions where conventional monitoring methods reach their limits, but which play a vital role in global ecosystem services.

## Acknowledgements

We thank the Fundación Jocotoco and Fundación Tesoro Escondido (Citlalli Morelos-Juarez) for logistic support and permission to do research on their reserves. We especially acknowledge local support from the staff: Katrin Krauth (manager of the Chocó Lab); Bryan Tamayo (plot manager); Lady Condoy, Leonardo de la Cruz, Franklin Quintero, Jefferson Tacuri, Jordy Ninabanda, Sílvia Vélez, Ismael Castellano, Fredi Cedeño (parabiologists); Alcides Zambrano (Canandé reserve staff); Citlalli Morelos-Juarez, Yadira Giler, Patricio Encarnacion, Ariel Villigu, Patricio Paredes and Adriana Argoti (Tesoro Escondido reserve staff). We thank the Ministry of Environment of Ecuador for granting research and collection permits through Contrato Marco MAE-DNB-CM-2021-0187. SK acknowledges funding by the Deutsche Forschungsgemeinschaft (BETA-FOR: Enhancing Structural Diversity in Production Forests (FOR 5375)); MK acknowledges funding by the Bavarian Research Institute for Digital Transformation (bidt), an institute of the Bavarian Academy of Sciences and Humanities (ROOT: Real-time earth Observation of fOrest dynamics and biodiversiTy (KON-22-024)).

## Supporting Information

**Supplementary Figure S1.**
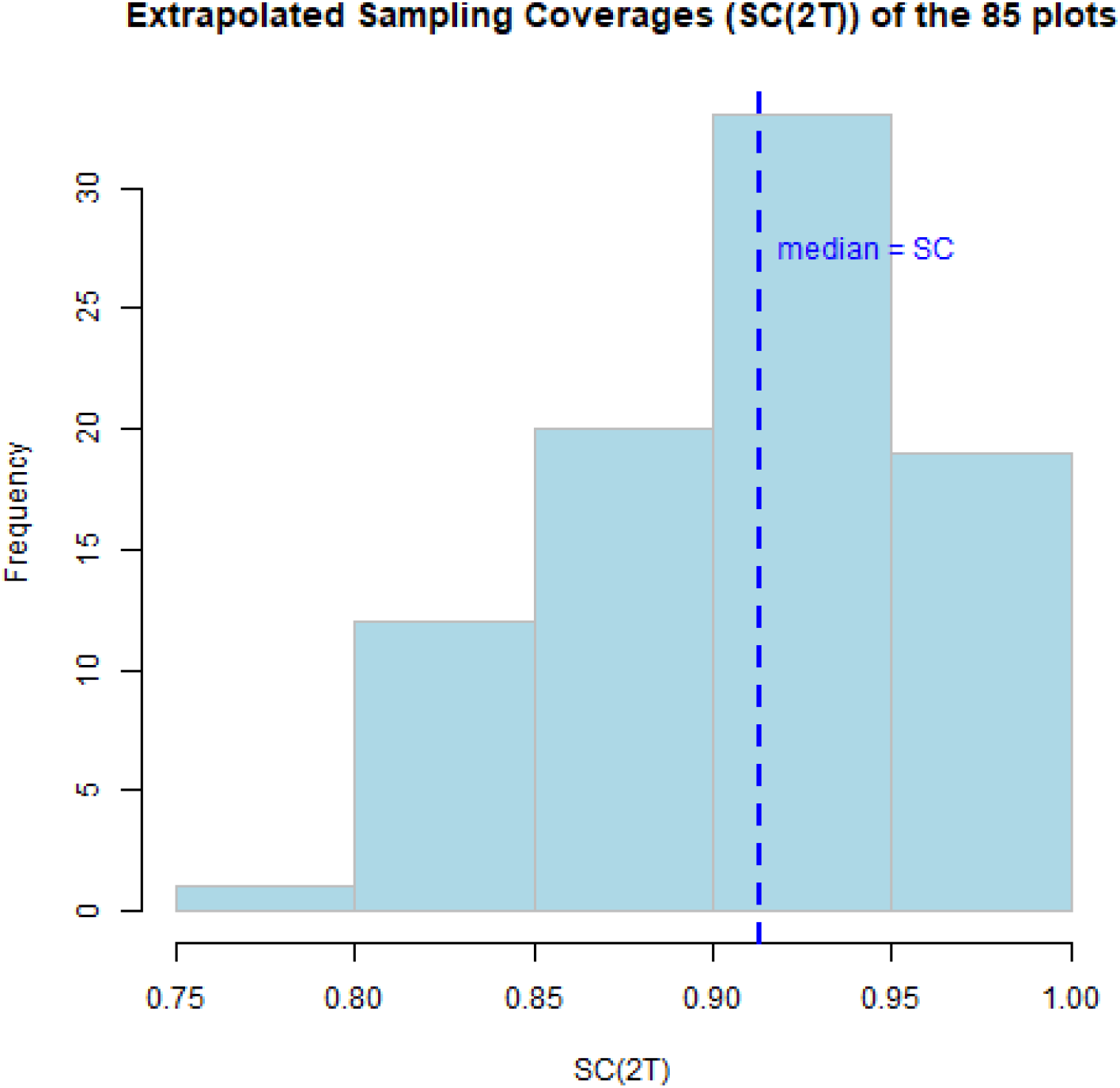
shows the distribution of the extrapolated sampling coverages across our 85 study plots. The sampling coverages represent the completeness of the acoustic data collected in each plot. The blue dashed line indicates the median sampling coverage value (0.913), which was used as the representative coverage level for the analysis.

**Supplementary Figure S2.**
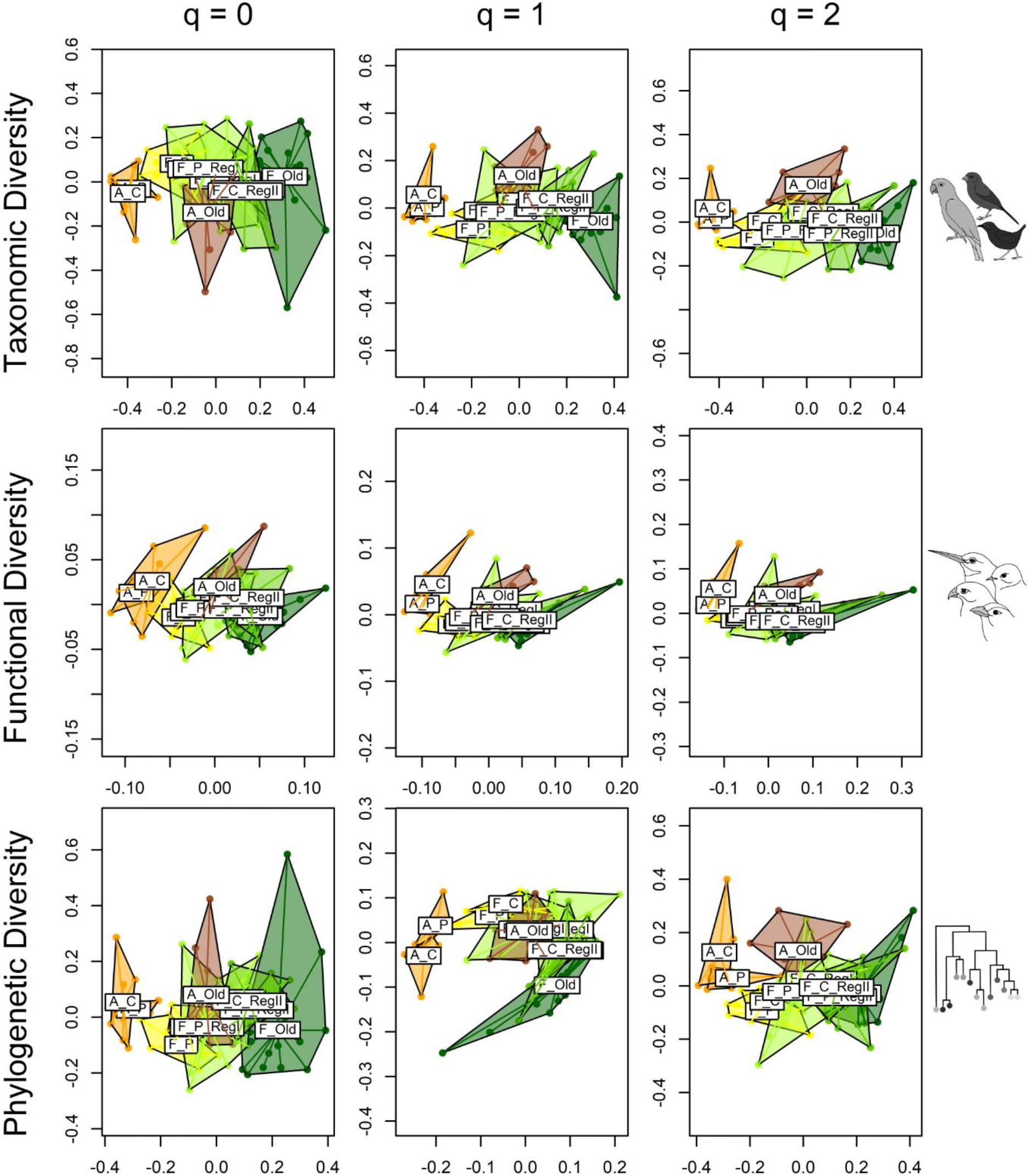
plots the first two nmds axes (NMDS1, NMDS2). The plots are arranged by diversity type (rows: Taxonomic, Functional, Phylogenetic) and orders of q (columns: q = 0, q = 1, q = 2). The category labels consist of up to three components: first, the matrix type (“A” for “agricultural matrix”, “F” for “forest matrix”), second, the type of (previous) land use (“Caca” or “C” for cacao plantation, “Past” or “P” for pasture) and third, the regeneration level: “Reg1” for 1-20 years since active use, and “Reg2” for 21-38 years of regeneration. “Old” stands for old-growths forests. The color palette of the “forest matrix” plots ranges from yellow to dark green, the “agricultural matrix” plots are colored in shades of orange and brown.

**Supplementary Figure S3.**
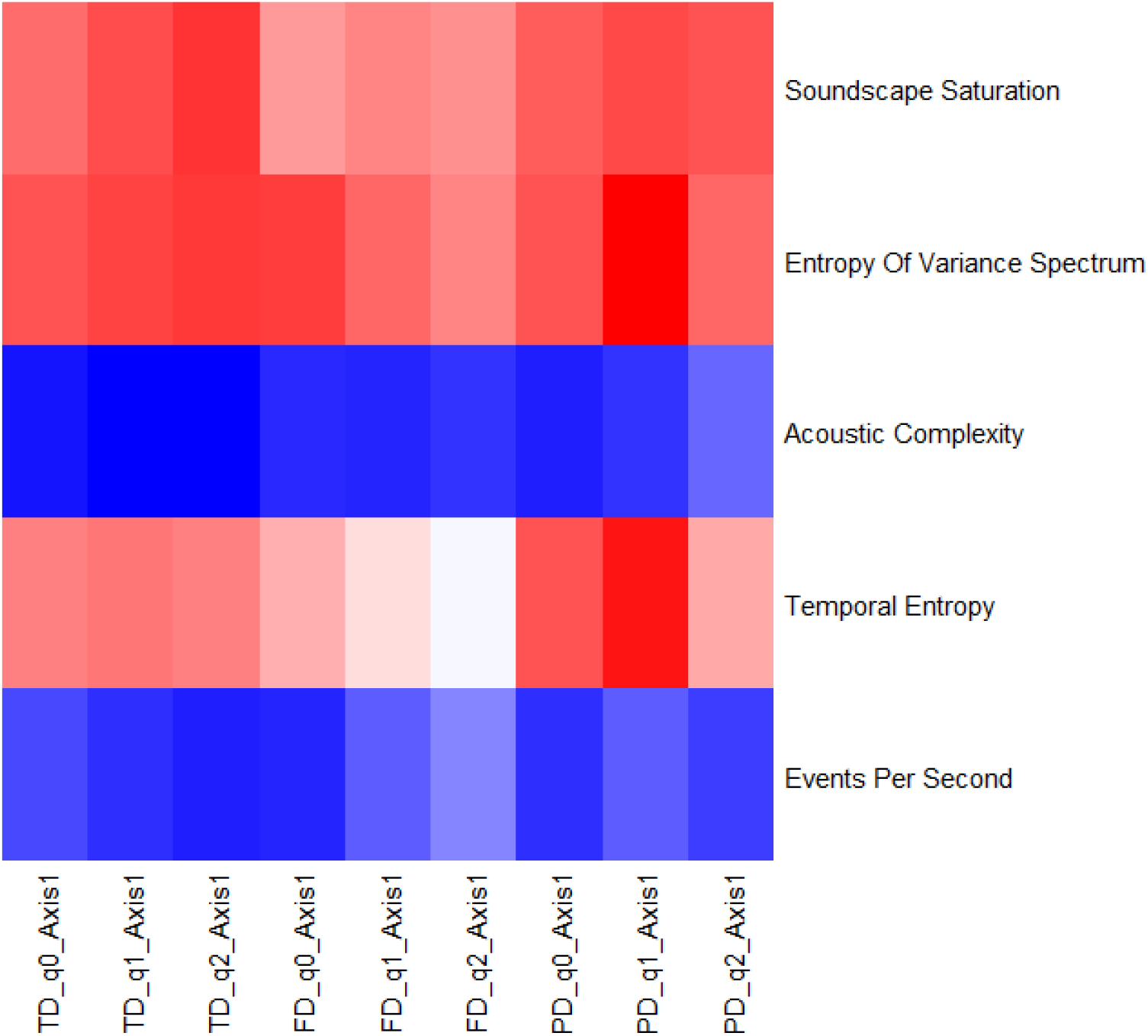
is a heatmap representation of (Table 1). The (intensity of the) t-values of the five acoustic indices (Soundscape Saturation, Entropy of Variance Spectrum, Acoustic Complexity, Temporal Entropy, and Events per Second) for predicting NMDS axis 1 across the three diversity types (Taxonomic, Functional and Phylogenetic) and orders of q (q = 0, q = 1, q = 2) are represented as colors, ranging from blue (negative t-values), through white (near-zero t-values) to red (positive t-values).

## REFERENCES

Alcocer, Irene; Lima, Herlander; Sugai, Larissa Sayuri Moreira; Llusia, Diego (2022): Acoustic indices as proxies for biodiversity: a meta-analysis. In: Biological reviews of the Cambridge Philosophical Society 97 (6), S. 2209–2236. DOI: 10.1111/brv.12890.

Antonelli, Alexandre; Rueda, Ximena; Calcagno, Robert; Nantongo Kalunda Pauline (2024): How biodiversity credits could help to conserve and restore nature. In: Nature 634 (8036), S. 1045–1049. DOI: 10.1038/d41586-024-03475-2.

Asner, Gregory P.; Martin, Roberta E.; Carranza-Jiménez, Loreli; Sinca, Felipe; Tupayachi, Raul; Anderson, Christopher B.; Martinez, Paola (2014): Functional and biological diversity of foliar spectra in tree canopies throughout the Andes to Amazon region. In: The New phytologist 204 (1), S. 127–139. DOI: 10.1111/nph.12895.

Borker, Abraham L.; Buxton, Rachel T.; Jones, Ian L.; Major, Heather L.; Williams, Jeffrey C.; Tershy, Bernie R.; Croll, Donald A. (2020): Do soundscape indices predict landscape-scale restoration outcomes? A comparative study of restored seabird island soundscapes. In: Restoration Ecology 28 (1), S. 252–260. DOI: 10.1111/rec.13038.

Bregman, Tom P.; Lees, Alexander C.; MacGregor, Hannah E. A.; Darski, Bianca; Moura, Nárgila G. de; Aleixo, Alexandre et al. (2016): Using avian functional traits to assess the impact of land-cover change on ecosystem processes linked to resilience in tropical forests. In: Proceedings. Biological sciences 283 (1844). DOI: 10.1098/rspb.2016.1289.

Burivalova, Zuzana; Towsey, Michael; Boucher, Tim; Truskinger, Anthony; Apelis, Cosmas; Roe, Paul; Game, Edward T. (2018): Using soundscapes to detect variable degrees of human influence on tropical forests in Papua New Guinea. In: Conservation Biology 32 (1), S. 205–215. DOI: 10.1111/cobi.12968.

Cadotte, Marc W.; Dinnage, Russell; Tilman, David (2012): Phylogenetic diversity promotes ecosystem stability. In: Ecology 93 (sp8). DOI: 10.1890/11-0426.1.

Chao, Anne; Chiu, Chun-Huo; Jost, Lou (2014): Unifying Species Diversity, Phylogenetic Diversity, Functional Diversity, and Related Similarity and Differentiation Measures Through Hill Numbers. In: Annu. Rev. Ecol. Evol. Syst. 45 (1), S. 297–324. DOI: 10.1146/annurev-ecolsys-120213-091540.

Chao, Anne; Jost, Lou (2012): Coverage-based rarefaction and extrapolation: standardizing samples by completeness rather than size. In: Ecology 93 (12), S. 2533–2547. DOI: 10.1890/11-1952.1.

Chazdon, Robin L.; Peres, Carlos A.; Dent, Daisy; Sheil, Douglas; Lugo, Ariel E.; Lamb, David et al. (2009): The potential for species conservation in tropical secondary forests. In: Conservation biology : the journal of the Society for Conservation Biology 23 (6), S. 1406–1417. DOI: 10.1111/j.1523-1739.2009.01338.x.

Díaz, Sandra; Settele, Josef; Brondízio, Eduardo S.; Ngo, Hien T.; Agard, John; Arneth, Almut et al. (2019): Pervasive human-driven decline of life on Earth points to the need for transformative change. In: Science (New York, N.Y.) 366 (6471). DOI: 10.1126/science.aax3100.

Dixon, Philip (2003): VEGAN, a package of R functions for community ecology. In: J Vegetation Science 14 (6), S. 927–930. DOI: 10.1111/j.1654-1103.2003.tb02228.x.

Dröge, Saskia; Martin, Dominic Andreas; Andriafanomezantsoa, Rouvah; Burivalova, Zuzana; Fulgence, Thio Rosin; Osen, Kristina et al. (2021): Listening to a changing landscape: Acoustic indices reflect bird species richness and plot-scale vegetation structure across different land-use types in north-eastern Madagascar. In: Ecological Indicators 120, S. 106929. DOI: 10.1016/j.ecolind.2020.106929.

Dunn, Robert R. (2004): Recovery of Faunal Communities During Tropical Forest Regeneration. In: Conservation Biology 18 (2), S. 302–309. DOI: 10.1111/j.1523-1739.2004.00151.x.

Edwards, David P.; Massam, Michael R.; Haugaasen, Torbjørn; Gilroy, James J. (2017): Tropical secondary forest regeneration conserves high levels of avian phylogenetic diversity. In: Biological Conservation 209, S. 432–439. DOI: 10.1016/j.biocon.2017.03.006.

Escobar, Sebastián; Newell, Felicity L.; Endara, María-José; Guevara-Andino, Juan E.; Landim, Anna R.; Neuschulz, Eike Lena et al. (2025): Reassembly of a tropical rainforest: A new chronosequence in the Chocó tested with the recovery of tree attributes. In: Ecosphere 16 (2), Artikel e70157. DOI: 10.1002/ecs2.70157.

Farina, Almo (2025): The acoustic complexity index (ACI): theoretical foundations, applied perspectives and semantics. In: Oikos 2025 (1), Artikel e10760. DOI: 10.1111/oik.10760.

Flowers, Colton; Le Tourneau, François-Michel; Merchant, Nirav; Heidorn, Brian; Ferriere, Régis; Harwood, Jake (2021): Looking for the-scape in the sound: Discriminating soundscapes categories in the Sonoran Desert using indices and clustering. In: Ecological Indicators 127, S. 107805. DOI: 10.1016/j.ecolind.2021.107805.

Ford, Helen V.; Schrodt, Franziska; Zieritz, Alexandra; Exton, Daniel A.; van der Heijden, Geertje; Teague, Jonathan et al. (2024): A technological biodiversity monitoring toolkit for biocredits. In: Journal of Applied Ecology 61 (9), S. 2007–2019. DOI: 10.1111/1365-2664.14725.

Gibb, Rory; Browning, Ella; Glover-Kapfer, Paul; Jones, Kate E. (2019): Emerging opportunities and challenges for passive acoustics in ecological assessment and monitoring. In: Methods Ecol Evol 10 (2), S. 169–185. DOI: 10.1111/2041-210X.13101.

Gindhart, Rosa; Müller, Jörg; Burivalova, Zuzana; Blüthgen, Nico; Busse, Annika; La Hoz, Maria de et al. (2024): The impact of land use on the acoustic behaviour of cicadas in the Chocó lowland tropical forest of Ecuador. In: Insect Conserv Diversity, Artikel icad.12793. DOI: 10.1111/icad.12793.

Hill, M. O. (1973): Diversity and Evenness: A Unifying Notation and Its Consequences. In: Ecology 54 (2), S. 427–432. DOI: 10.2307/1934352.

Hubau, Wannes; Lewis, Simon L.; Phillips, Oliver L.; Affum-Baffoe, Kofi; Beeckman, Hans; Cuní-Sanchez, Aida et al. (2020): Asynchronous carbon sink saturation in African and Amazonian tropical forests. In: Nature 579 (7797), S. 80–87. DOI: 10.1038/s41586-020-2035-0.

Hughes, Emma C.; Edwards, David P.; Sayer, Catherine A.; Martin, Philip A.; Thomas, Gavin H. (2020): The effects of tropical secondary forest regeneration on avian phylogenetic diversity. In: Journal of Applied Ecology 57 (7), S. 1351–1362. DOI: 10.1111/1365-2664.13639.

Isbell, Forest; Gonzalez, Andrew; Loreau, Michel; Cowles, Jane; Díaz, Sandra; Hector, Andy et al. (2017): Linking the influence and dependence of people on biodiversity across scales. In: Nature 546 (7656), S. 65–72. DOI: 10.1038/nature22899.

Jetz, W.; Thomas, G. H.; Joy, J. B.; Hartmann, K.; Mooers, A. O. (2012): The global diversity of birds in space and time. In: Nature 491 (7424), S. 444–448. DOI: 10.1038/nature11631.

Jin, Yi; Qian, Hong (2023): U.PhyloMaker: An R package that can generate large phylogenetic trees for plants and animals. In: Plant diversity 45 (3), S. 347–352. DOI: 10.1016/j.pld.2022.12.007.

Kortmann, Mareike; Chao, Anne; Schaefer, H. Martin; Blüthgen, Nico; Gelis, Rudy; Tremlett, Constance J. et al. (2025): Sample coverage affects diversity measures of bird communities along a natural recovery gradient of abandoned agriculture in tropical lowland forests. In: Journal of Applied Ecology, Artikel 1365–2664.14879. DOI: 10.1111/1365-2664.14879.

Lamont, Timothy A. C.; Williams, Ben; Chapuis, Lucille; Prasetya, Mochyudho E.; Seraphim, Marie J.; Harding, Harry R. et al. (2022): The sound of recovery: Coral reef restoration success is detectable in the soundscape. In: Journal of Applied Ecology 59 (3), S. 742–756. DOI: 10.1111/1365-2664.14089.

Le Toan, T.; Quegan, S.; Davidson, M.W.J.; Balzter, H.; Paillou, P.; Papathanassiou, K. et al. (2011): The BIOMASS mission: Mapping global forest biomass to better understand the terrestrial carbon cycle. In: Remote Sensing of Environment 115 (11), S. 2850–2860. DOI: 10.1016/j.rse.2011.03.020.

Mouillot, David; Bellwood, David R.; Baraloto, Christopher; Chave, Jerome; Galzin, Rene; Harmelin-Vivien, Mireille et al. (2013): Rare species support vulnerable functions in high-diversity ecosystems. In: PLoS biology 11 (5), e1001569. DOI: 10.1371/journal.pbio.1001569.

Müller, Jörg; Mitesser, Oliver; Schaefer, H. Martin; Seibold, Sebastian; Busse, Annika; Kriegel, Peter et al. (2023): Soundscapes and deep learning enable tracking biodiversity recovery in tropical forests. In: Nature communications 14 (1), S. 6191. DOI: 10.1038/s41467-023-41693-w.

Pillay, Rajeev; Venter, Michelle; Aragon-Osejo, Jose; González-Del-Pliego, Pamela; Hansen, Andrew J.; Watson, James Em; Venter, Oscar (2022): Tropical forests are home to over half of the world’s vertebrate species. In: Frontiers in ecology and the environment 20 (1), S. 10–15. DOI: 10.1002/fee.2420.

Prevedello, Jayme Augusto; Vieira, Marcus Vinícius (2010): Does the type of matrix matter? A quantitative review of the evidence. In: Biodivers Conserv 19 (5), S. 1205–1223. DOI: 10.1007/s10531-009-9750-z.

Rozendaal, Danaë M. A.; Bongers, Frans; Aide, T. Mitchell; Alvarez-Dávila, Esteban; Ascarrunz, Nataly; Balvanera, Patricia et al. (2019): Biodiversity recovery of Neotropical secondary forests. In: Science advances 5 (3), eaau3114. DOI: 10.1126/sciadv.aau3114.

Sun, Yuren; Midori Maeda, Tatiana; Solís-Lemus, Claudia; Pimentel-Alarcón, Daniel; Buřivalová, Zuzana (2022): Classification of animal sounds in a hyperdiverse rainforest using convolutional neural networks with data augmentation. In: Ecological Indicators 145, S. 109621. DOI: 10.1016/j.ecolind.2022.109621.

Teixeira, Daniella; Maron, Martine; van Rensburg, Berndt J. (2019): Bioacoustic monitoring of animal vocal behavior for conservation. In: Conservat Sci and Prac 1 (8), Artikel e72. DOI: 10.1111/csp2.72.

Thompson, Michelle E.; Donnelly, Maureen A. (2018): Effects of Secondary Forest Succession on Amphibians and Reptiles: A Review and Meta-analysis. In: Copeia 106 (1), S. 10–19. DOI: 10.1643/CH-17-654.

Thorn, Simon; Bässler, Claus; Brandl, Roland; Burton, Philip J.; Cahall, Rebecca; Campbell, John L. et al. (2018): Impacts of salvage logging on biodiversity: a meta-analysis. In: Journal of Applied Ecology 55 (1), S. 279–289. DOI: 10.1111/1365-2664.12945.

Tobias, Joseph A.; Sheard, Catherine; Pigot, Alex L.; Devenish, Adam J. M.; Yang, Jingyi; Sayol, Ferran et al. (2022): AVONET: morphological, ecological and geographical data for all birds. In: Ecology letters 25 (3), S. 581–597. DOI: 10.1111/ele.13898.

Vardon, Michael J.; Lindenmayer, David B. (2023): Biodiversity market doublespeak. In: Science (New York, N.Y.) 382 (6670), S. 491. DOI: 10.1126/science.adg6823.

Vega-Hidalgo, Álvaro; Flatt, Eleanor; Whitworth, Andrew; Symes, Laurel (2021): Acoustic assessment of experimental reforestation in a Costa Rican rainforest. In: Ecological Indicators 133, S. 108413. DOI: 10.1016/j.ecolind.2021.108413.

Xie, Jiangjian; Zhong, Yujie; Zhang, Junguo; Liu, Shuo; Ding, Changqing; Triantafyllopoulos, Andreas (2023): A review of automatic recognition technology for bird vocalizations in the deep learning era. In: Ecological Informatics 73, S. 101927. DOI: 10.1016/j.ecoinf.2022.101927.

Zamora-Gutierrez, Veronica; Lopez-Gonzalez, Celia; MacSwiney Gonzalez, M. Cristina; Fenton, Brock; Jones, Gareth; Kalko, Elisabeth K. V. et al. (2016): Acoustic identification of Mexican bats based on taxonomic and ecological constraints on call design. In: Methods Ecol Evol 7 (9), S. 1082–1091. DOI: 10.1111/2041-210X.12556.

